# Inhibition of DKK-1 by WAY262611 Inhibits Osteosarcoma Metastasis

**DOI:** 10.1101/2024.12.10.627181

**Authors:** Adit Tal, Shimara Gunawardana-Zeigler, Da Peng, Yuqi Tan, Natalia Munoz Perez, Rachel Offenbacher, Laurel Kastner, Paul Ciero, Matthew E. Randolph, Yun Gong, Hong-Wen Deng, Patrick Cahan, David M. Loeb

## Abstract

Osteosarcoma (OS) is the most common primary malignant bone tumor in childhood. Patients who present with metastatic disease at diagnosis or relapse have a very poor prognosis, and this has not changed over the past four decades. The Wnt signaling pathway plays a role in regulating osteogenesis and is implicated in OS pathogenesis. DKK-1 inhibits the canonical Wnt signaling pathway, causing inhibition of osteoblast differentiation and disordered bone repair. Our lab previously demonstrated that a monoclonal antibody against DKK-1 prevented metastatic disease in a mouse model. This study expands upon those findings by demonstrating similar results with a small molecule inhibitor of DKK-1, WAY262611, both *in vitro* and *in vivo*. WAY262611 was evaluated *in vitro* on osteosarcoma cell lines, including proliferation, caspase activation, cell cycle analysis, and signaling pathway activation. We utilized our orthotopic implantation-amputation model of osteosarcoma metastasis *in vivo* to determine the impact of WAY262611 on primary tumor progression and metastatic outgrowth of disseminated tumor cells. Differentiation status was determined using single cell RNA sequencing. We show here that WAY262611 activates canonical Wnt signaling, enhances nuclear localization and transcriptional activity of beta-catenin, and slows proliferation of OS cell lines. We also show that WAY262611 induces osteoblastic differentiation of an OS patient-derived xenograft *in vivo*, as well as inhibiting metastasis. This work credentials DKK-1 as a therapeutic target in OS, allowing for manipulation of the Wnt signaling pathway and providing preclinical justification for the development of new biologics for prevention of osteosarcoma metastasis.

## Introduction

Osteosarcoma (OS) is the most common primary malignant bone tumor in childhood.[1] Surgery alone cures only a small minority of patients, but since the introduction of systemic chemotherapy, long-term survival of patients with localized, non-metastatic disease now approaches 75%.[2] Despite this achievement, patients with metastatic disease at presentation have a significantly worse prognosis, with overall survival of 20-30%, and there has been little improvement in the last forty years despite maximizing the intensity of chemotherapy[3] or adjusting treatment based on tumor response to therapy.[4, 5] Additionally, 25% of patients initially diagnosed with localized tumors will suffer a metastatic relapse and succumb to their disease.[6] Optimization of treatment for children with metastatic and relapsed OS will require novel approaches to therapy based on understanding the pathophysiology of metastatic spread, including identifying and targeting genomic aberrations and activated signaling pathways that drive this process. Although novel therapeutic targets have been investigated in clinical trials, including microtubule inhibitors, immune modulators, and monoclonal antibodies to GD2, VEGF, and PD-1 [7], none of these are specific to metastatic spread, and none have shown efficacy in preventing or treating OS metastasis. To date, there have been no agents specifically designed to target the process of metastasis; therefore, our work is focused on identifying and targeting biological pathways that are critical to the process of metastasis, rather than focusing on developing new agents to shrink existing tumors.

The Wnt signaling pathway is crucial to embryonic development and tissue homeostasis, plays a role in regulating osteogenic differentiation, and has been studied in the context of osteosarcoma progression.[8] The Wnt family comprises 19 secreted glycoproteins, which bind to transmembrane Frizzled receptors or tyrosine kinases that activate various intracellular signaling pathways: a canonical, beta catenin-dependent pathway and several noncanonical, beta catenin-independent pathways.[9] The canonical pathway is central to osteoblast formation, as previous studies have shown that increased beta catenin is found in cells committed to the osteoblast lineage. The dickkopf family of proteins, which interact directly with LRP5/6 and Wnt receptors to inhibit canonical signaling, are negative regulators of this pathway. One such protein is DKK-1, which counteracts the Wnt signaling-mediated effects on bone homeostasis, causing inhibition of osteoblast differentiation, osteoclast formation, increased bone resorption, and disordered bone repair.[10]

The role of Wnt signaling in the pathogenesis of osteosarcoma is controversial; it has been implicated in both supporting the proliferation of bone sarcoma cells and in driving healthy osteogenic differentiation and preventing further tumorigenesis. DKK-1 levels are significantly elevated in cancers with osteolytic metastases, such as prostate cancer and breast cancer [11, 12], suggesting that bone breakdown can create a tumor microenvironment causing disarray to the normal bone niche and permitting tumor expansion and metastasis. Human OS cell lines express DKK-1 mRNA, and the protein is actively secreted by rapidly proliferating OS cells, suggesting that activation of the canonical Wnt signaling pathway might be anti-tumorigenic.[13] Congruous to this, the Gregory lab found elevated levels of DKK-1 in the plasma of pediatric patients with OS compared to healthy controls. They demonstrated that DKK-1 is expressed maximally at the periphery of OS tumors *in vivo* and in rapidly proliferating cells *in vitro*, and that DKK-1 levels reflected the number of surviving OS cells in *in vivo* tumor models.[14] We hypothesize that inhibition of DKK-1 could be a novel approach to prevent OS metastasis and relapse. We have previously demonstrated that a neutralizing monoclonal antibody against DKK-1 slows tumor growth, inhibits metastasis, and induces markers of differentiation in OS patient-derived xenografts (PDX) implanted in the tibias of immune deficient mice.[15] The production of that monoclonal antibody was unfortunately halted and further access was denied by the pharmaceutical company. Thus, we now describe the use of a small molecule inhibitor of DKK-1, designated WAY262611, as an alternative way to augment canonical Wnt signaling in OS cells and evaluated its effect on tumor growth and metastasis *in vivo*. This work provides additional support to the concept of inhibiting DKK-1 to drive OS differentiation and prevent metastasis, credentialing DKK-1 as an important therapeutic target.

## Materials and Methods

### Cell Lines and Xenografts

3 osteosarcoma cell lines, U2OS (CVCL 0042), HOS (CVCL 0312), and SaOS2 (CVCL 0548), originally provided by ATCC and through a generous gift from the Hoang Lab (AECOM), were maintained in RPMI 1640 media supplemented with 10% FBS. Cells were incubated in a ThermoScientific HERACell Vios 160i CO_2_ incubator at 37 °C. Cells were split every 3 days and incubated at different confluences per experiment. Cell lines were authenticated using short tandem repeat (STR) profiling. Periodic assessment for mycoplasma contamination was performed using the Lonza MycoAlert Mycoplasma Detection Kit. The DAR OS PDX, created from cells isolated from a malignant pleural effusion in an osteosarcoma patient, was a generous gift from Dr. Chand Khanna (NCI, NIH).

### DKK-1 small molecule inhibitor

WAY262611 (catalog #17704) was purchased from Cayman Chemical Company. For *in vitro* experiments, we determined the sensitivity of OS cell lines to WAY262611 using a CCK8 proliferation assay. For *in vivo* experiments, drug was diluted in DMSO at a concentration of 30 mg/mL. For *in vitro* experiments, drug was diluted in ethanol at a concentration of 25 mg/mL.

### *In Vitro* Incucyte Proliferation and Caspase Activation Assays

To determine the IC50 for proliferation, 2,500 U2OS, SaOS2 and HOS cells were plated per well in a 96 well, tissue culture-treated plate (Corning) with 75 uL of RPMI 1640 (Gibco) + 10% FBS and maintained under standard conditions. After 24 hours to allow for adherence, WAY262611 was added at various concentrations from 0.05 µM to 10 mM. Plates were inserted into the Incucyte^®^Live-Cell Analysis System and confluence was measured utilizing live-cell time-lapse imaging.

To determine the cytotoxic effect of WAY262611 on osteosarcoma cell lines, 2,500 U2OS, SaOS2 and HOS cells were plated per well in a 96 well, tissue culture-treated plate with 75 uL of RPMI 1640 + 10% FBS and maintained under standard conditions. After 24 hours to allow for adherence, media was refreshed, Caspase 3/7 dye was added, and WAY26211 treatment was initiated at the indicated concentrations. Plates were inserted into the Incucyte® Live-Cell Analysis System and caspase activity was measured utilizing live-cell time-lapse imaging over 72 hours.

### Cell cycle analysis by flow cytometry

Five hundred thousand U2OS, SaOS2, and HOS cells were plated per well in a 6 well, tissue culture-treated plate with 3 mL of RPMI 1640 + 10% FBS and maintained under standard conditions. After 24 hours to allow for adherence, WAY262611 was added at the previously determined IC50. After 36 hours of treatment, flow cytometry was performed utilizing Click-iT™ EdU Cell Proliferation Kit for Imaging, Alexa Fluor™ 647 dye and FxCycle™ Violet stain (F10347).

### Protein Analysis and Western blotting for DKK-1 Quantification

Protein was isolated from OS cell lines using the Santa Cruz RIPA lysis buffer system and quantified by BCA. Western blots were run on the BioRad Mini Protean TetraCell and transferred using the BioRad Trans-Blot Turbo transfer system. Antibodies to DKK-1 were obtained from Abcam (Catalog #61034) and Origene (Catalog #TA349295). Chemiluminescence imaging was performed on a BioRad ChemiDoc Touch imaging system.

### Differentiation Assay

Differentiation assays were performed using osteogenic medium according to the Bio-Protocol “Osteogenic and Adipogenic Differentiation of Osteosarcoma Cells”[16] with or without WAY262611. RNA was collected from cells on Day 0, after exposure to drug in regular growth medium, and then on Day 6 after 6 days in differentiation medium with or without WAY26261 for further analysis by qRT-PCR for quantification of osteogenic markers.

### RNA Isolation and qRT-PCR

RNA was isolated from OS cell lines after differentiation assays using a Qiagen RNAeasy Plus Mini Kit and cDNA was synthesized using the BioRad iScript RT Supermix. Primers for osteogenic differentiation markers were obtained from BioRad (Catalog #10025636): ALPL (Unique assay ID qHsaCED0045991) and Sino Biological (SPP-1 HP100389). Quantitative RT-PCR was then performed using the Applied Biosystems QuantStudio5 Real-Time PCR thermal cycler. The cycle threshold (C_t_) values were calculated using the 2^-ΔΔCt^ method, and qRT-PCR and melting curve analysis were conducted using the ThermoFisher data analysis cloud.

### Immunofluorescence

Cultured cells were seeded on six-well plates at a density of 2.5 x 10^5^ cells per well, on coverslips until 50% confluence overnight in an incubator at 37 °C, fixed with 4% paraformaldehyde, washed with PBS, and permeabilized with 0.2% Triton X. Cells were then blocked with 5% goat serum, 2% BSA, and 0.2% Triton X in PBS for 1 hour and incubated in blocking media with primary antibody, or isotype control, in a humidified chamber for 1 hour at room temperature. Cells were subsequently washed and incubated for 1 hour in a humidified chamber with secondary antibody and DAPI, and then coverslips were mounted on slides. Coverslips and slides were placed to dry overnight at 4 °C in an enclosed slide box, and were imaged within 5 days. Primary antibody to beta catenin was obtained from BD Transduction Laboratories (Catalog #610153). Slides were imaged on the Leica SP5 confocal microscope at 40x and 63X magnification. Nuclear:cytoplasmic calculation and image analysis was performed with the help of the Analytical Imaging Facility and using Volocity software by Quorum Technologies.

### Stable Luciferase Assay

To make stable luciferase cell lines, 7TCF-Firefly luciferase (Addgene 24308) and pLX313-Renilla luciferase (Addgene 24308) lentivirus constructs were obtained from Addgene. Lentivirus was produced by transfecting HEK293T cells (CVCL 0063) with 7TCF-Firefly luciferase or pLX313-Renilla luciferase constructs for 48 hours. Osteosarcoma cell lines (U2OS, HOS, and SaOS2) were first transduced with lentivirus packaged with 7TCF-Firefly luciferase overnight and selected for 72 hours with puromycin. Positively infected osteosarcoma cells were subsequently transduced with lentivirus packaged with pLX313-Renilla luciferase and selected with hygromycin for 72 hours. Luciferase activity was assayed using the Promega Dual-Luciferase Reporter Assay System. A tankyrase inhibitor, JW-55 (Tocris, Catalog #4514) was also used for these experiments.

### CRISPR/Cas9 mediated DKK-1 KO

DKK-1 Knockout CRISPR kit was purchased from Origene (Catalog #KN504599), which included a linear donor tagged with green fluorescent protein (GFP) and puromycin and 2 different target sequences (gRNA1 and gRNA2). Twenty-four hours prior to transfection, cells were plated in 6 well plates at 2×10^5^ into 2 ml of culture media to obtain 50% confluence on the following day. Separate transfections were set up for each gRNA vector. Turbofectin reagent (Catalog #TF81001) was used for transfection as per Origene protocol, using 1 ug of gRNA vectors and 1 ug of donor DNA. Cells were incubated for 48 hours in culture media, split, and then puromycin selection media was used for additional passages. After 1 week of puromycin selection, cells were sorted by flow cytometry to collect cells with the strongest expression of GFP. Once the “high GFP” cohort was collected, monoclonal cell expansion was performed to isolate clones for knockout analysis. Thirty clones were cultured, and western blot assay was performed for assessment of knockout. Clones that were transfected but did not have a knockout were used as controls for experiments involving knockout cells.

### Animal studies

All NOD.Cg-Prkdc^scid^ Il2rg^tm1Wjl^/SzJ (IMSR JAX:005557) mice were maintained according to IACUC approved protocols in accordance with Albert Einstein College of Medicine research guidelines. Human derived tumor cells were xenografted into NSG “donor” mice subcutaneously, and were grown in order to provide enough tumor tissue for cohorted experiments. Fragments of tumor harvested from these “donor” mice were surgically implanted into the tibia of experimental mice. Once the tumors reached 1.5 cm maximal diameter, the limbs were amputated and the mice followed for recurrence or metastasis. Mice were euthanized when body-condition score (BCS) was found to be 2 or lower. Rate of primary tumor growth, time to amputation, metastatic index, and survival were statistically analyzed using GraphPad software (SCR 002798) via t-test, ANOVA, and the Cox proportional hazards model to account for several risk factors.

### Single-Cell Analysis RNA Sequencing

A freshly growing PDX, designated DAR was collected from a so-called “donor mouse” and rinsed in cold 1X DPBS. It was cut into 2-3 mm fragments and mixed with tissue digestion media (10 mL/g of tumor) containing RPMI 1640 without FBS, 0.1 mg/mL DNase I (Roche, Catalog#10104159001), and 0.125 mg/mL Liberase (Roche, Catalog#05401020001). The tumor-media mixture was incubated shaking (220 RPM) at 37°C for 20 minutes. Digestion was stopped by adding cold RPMI with 10% FBS and placing the mixture on ice for 7 minutes. The mixture then was passed through a 40 µm cell sieve to separate single cells from undigested tissue. The flow through was pelleted, incubated with ACK lysis buffer (Quality Biological, Catalog #118156101) to lyse red blood cells, and enriched for live cells using a dead cell removal kit (Miltenyi, Catalog#130090101). NGS libraries were prepared using Chromium Next GEM Single Cell 3’v3.1 (dual index) by 10x Genomics. Libraries with 1% Phix spike-in were sequenced at GENEWIZ using a NovaSeq 6000 Illumina sequencer with 2×150 bp reads. The raw fastq files were aligned to GRCh38 and mm10 combined transcriptome using 10x Genomics Cell Ranger version 7.0.1. [17] The cells that were categorized as mouse cells by Cell Ranger were discarded from downstream analysis. Standard scRNA-seq analysis was performed using Scanpy (version 1.9.5). The lowly expressed genes were first removed (min_cells = 100). The raw counts were log-normalized (target_sum = 1e4) and highly variable genes were selected. The gene expressions were scaled and Principal Component Analysis (PCA) was performed. The top 10 PCs were chosen to construct neighborhood graph for UMAP embeddings and Leiden clustering (resolution = 0.09).

Differential gene expression analysis was done using Scanpy’s ‘tl.rank_genes_groups’ function (default parameters) for each cell cluster.[18] Lowly expressed genes (less than 10% expression in target and background cell groups) were removed. The outputted scores from ‘tl.rank_genes_groups’ (z-scores) were used as gene ranks for gene set enrichment analysis. Gseapy (version 1.0.4) was used to perform gene set enrichment with pre-ranked correlation (permutation_num = 1000). [19]

To compare the osteosarcoma cells with normal human osteoblast cells, a pySingleCellNet [20] classifier was trained (parameters: nTopGenes = 100, nRand = 100, nTrees = 1000, nTopGenePairs = 100, stratify = True, limitToHVG = True) using normal human osteoblast cells from Gong et al. [21] All the scripts used to analyze the single cell expression data can be found on our Github page.

### Statistical Analysis

All statistical analyses in this study were conducted using GraphPad Prism 9 (GraphPad Software, San Diego, CA, USA) and all experiments were performed at least 3 times to confirm statistical significance and reproducibility. T-test was used for data comparison between two groups, and analysis of variance (ANOVA) was used for data comparison between three or more groups. Cox proportional hazards model was used for in vivo survival analysis. A p<0.05 was considered statistically significant.

### Data Availability

The data generated in this study are publicly available on Zenodo (record ID: 13914007) https://zenodo.org/records/13914007. We have placed all of our analysis code, provided detailed instructions to recreate the analysis and updated the links to download raw data on our GitHub repo https://github.com/CahanLab/DKK1_OS.

## Results

### WAY262611 slows the growth of OS cell lines in vitro

To validate the use of WAY262611 (chemical structure shown in Figure 1A), we investigated the level of expression of DKK-1 in OS cell lines. We demonstrated strong DKK-1 protein expression in all 3 OS cell lines by western blot analysis (Figure 1B). To begin to understand the impact of DKK-1 inhibition on osteosarcoma, we used a CCK8 assay and found that WAY262611 inhibits the viability of all 3 OS cell lines with an IC50 of 3-8 μM (Figure 1C). These results could reflect either an impact on proliferation or on cell death. To distinguish between these possibilities, we performed Incucyte proliferation assays. Using this approach, we found that WAY262611 inhibits the proliferation of OS cell lines with an IC50 of ∼1 µM in each cell line (Figure 2A-C). To confirm that WAY262611 is not cytotoxic, we performed caspase activity assays on OS cells. Using the Incucyte fluorescence-based assay of caspase activity, we found that WAY262611 does not significantly induce caspase 3 activity at concentrations that inhibit proliferation *in vitro* (Figure 2D-F). Finally, cell cycle analysis demonstrated that WAY262611 induces a G2/M cell cycle arrest at 3 uM in all three cell lines (Figure 2G-I). Thus, WAY262611 inhibits cell proliferation by inducing cell cycle arrest, and not through induction of apoptosis, at least *in vitro*.

**Figure 1.**
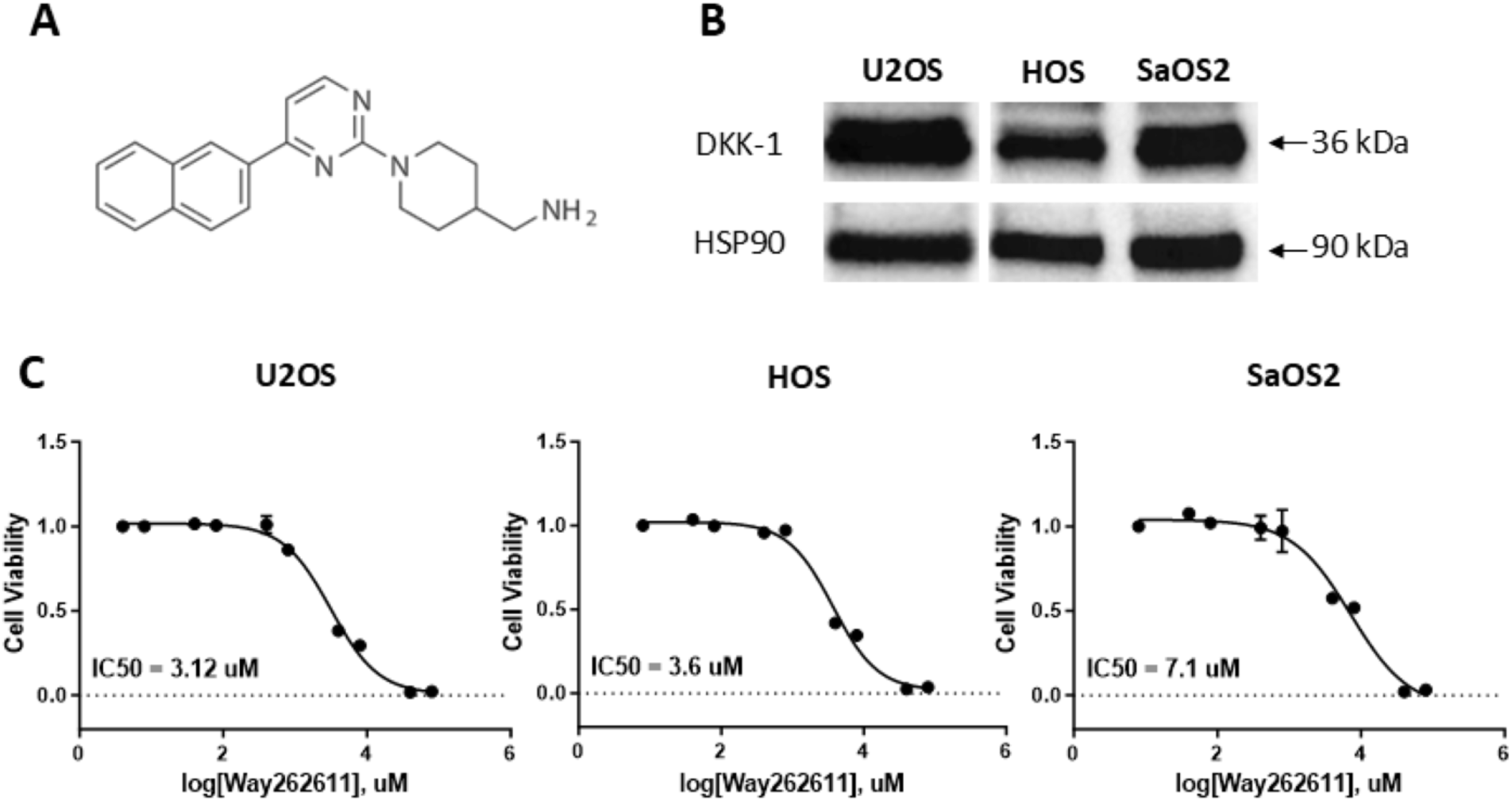
WAY262611 inhibits viability of osteosarcoma cells *in vitro*. **(A)** The chemical structure of WAY262611. (**B)** Western blot confirming expression of DKK-1 in U2OS, HOS, and SaOS2 cells. HSP90 is the loading control. (**C)** CCK8 viability assays showing dose-dependent decrease in viability of all 3 osteosarcoma cell lines when treated with WAY262611. Error bar is SEM of triplicate assays.

**Figure 2.**
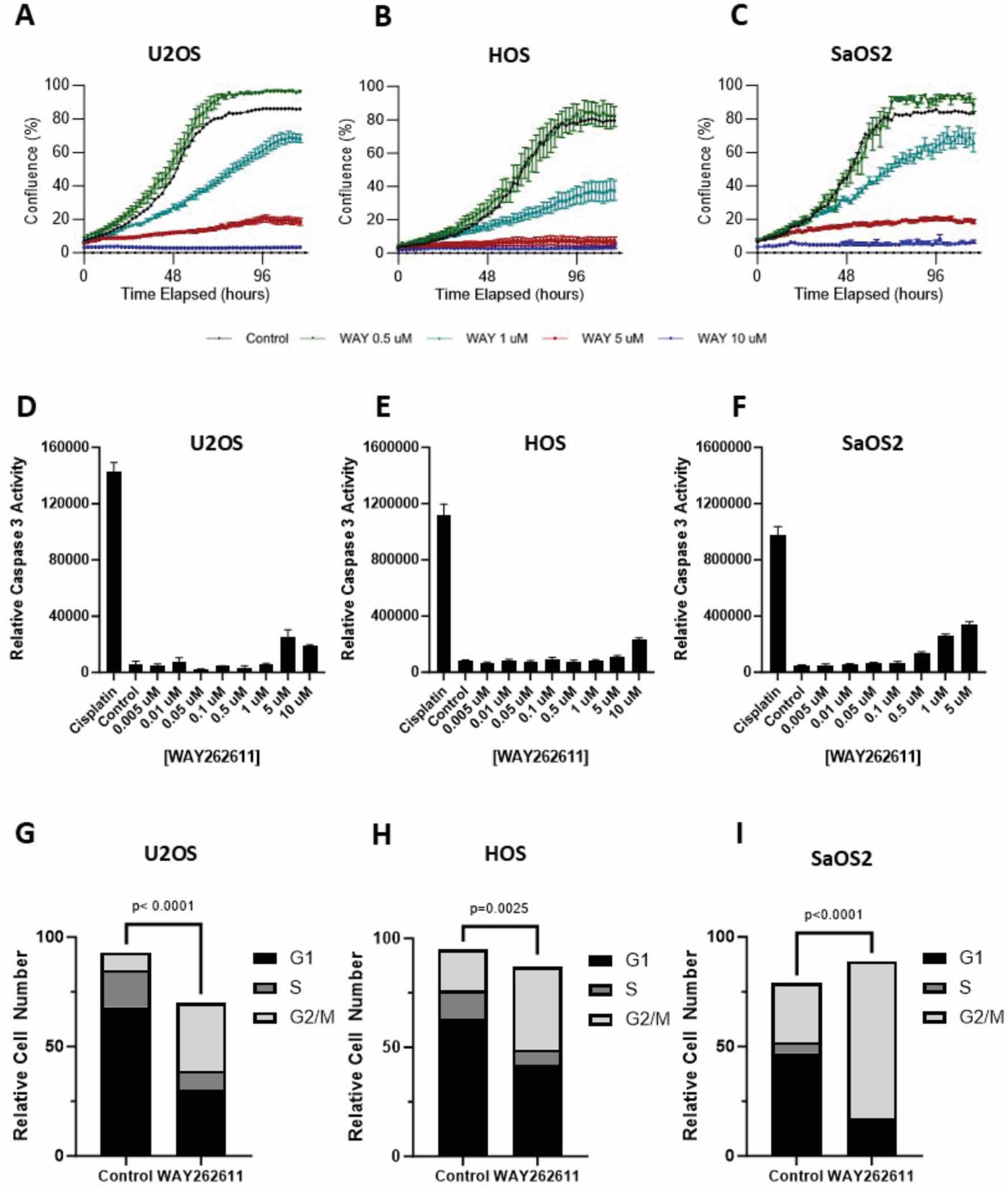
WAY262611 inhibits proliferation by inducing a G2/M cell cycle arrest. Treatment with WAY262611 induces a dose-dependent inhibition of proliferation in U2OS **(A)**, HOS **(B)**, and SaOS2 **(C)** cells. WAY262611 does not induce caspase 3 activity, even at very high doses, in U2OS **(D)**, HOS **(E)**, and SaOS2 **(F)** cells. Cell cycle analysis demonstrates a G2/M arrest in U2OS **(G)**, HOS **(H)**, and SaOS2 **(I)** cells treated with WAY262611 at the IC50. Differences between treated and untreated cells was statistically significant by chi-squared test. Error bar is the SEM of triplicate assays. All experiments were repeated at least 3 times.

### WAY262611 induces canonical Wnt signaling

To confirm that WAY262611-mediated inhibition of DKK-1 induces canonical Wnt signaling, we first investigated whether the drug induces nuclear translocation of beta catenin in OS cells. Based on the previously described IC50, cells were treated with 1, 3, 6 uM WAY262611 or vehicle control, and subcellular localization of beta catenin was evaluated by immunofluorescence. Maximum nuclear localization was identified at 6uM WAY262611 without significant cell toxicity (Figure 3D). In U2OS cells there was a 2-fold increase and in SaOS2 cells there was a 1.3-fold increase in beta catenin (green) fluorescence intensity in the nuclear compartment as compared to the intensity of green fluorescence within the total cellular compartment after treatment with WAY262611 for 24 hours. HOS cells demonstrated an earlier peak of intracellular beta catenin, a 1.7-fold increase at 5 hours (Figure 3A-C). Thus, in all 3 OS cell lines WAY262611 increased beta catenin nuclear localization, though with varying kinetics and strength of induction. Next, we used a functional assay to determine whether nuclear localization of beta catenin induced by WAY262611 causes increased beta catenin-driven transcription. OS cells were stably transfected with 7TCF-Firefly luciferase construct, a beta catenin reporter plasmid that contains the binding sites for Wnt transcription factors TCF driving expression of firefly luciferase. Cells were treated with 0.5-10 uM WAY262611 or vehicle control for 24 hours and then assayed for luciferase activity. Treatment with 6 uM WAY262611 induced a significant increase in luciferase activity in each cell line (Figure 3E-G). To confirm that the increase in luciferase activity is due to an increase in beta catenin-dependent Wnt signaling and not an off-target effect of the drug, we repeated the experiment in the presence or absence of the tankyrase inhibitor, JW-55. Tankyrase regulates the stability of the beta catenin destruction complex, so a tankyrase inhibitor should effectively prevent beta catenin-dependent signaling.[22] Indeed, we found that JW-55 inhibits the basal level of luciferase activity in each cell line, and also prevents induction of luciferase by WAY262611 (Figure 3E-G). These findings confirm that increased nuclear localization of beta catenin induced by WAY262611 results in the activation of canonical Wnt signaling in all 3 OS cell lines.

**Figure 3.**
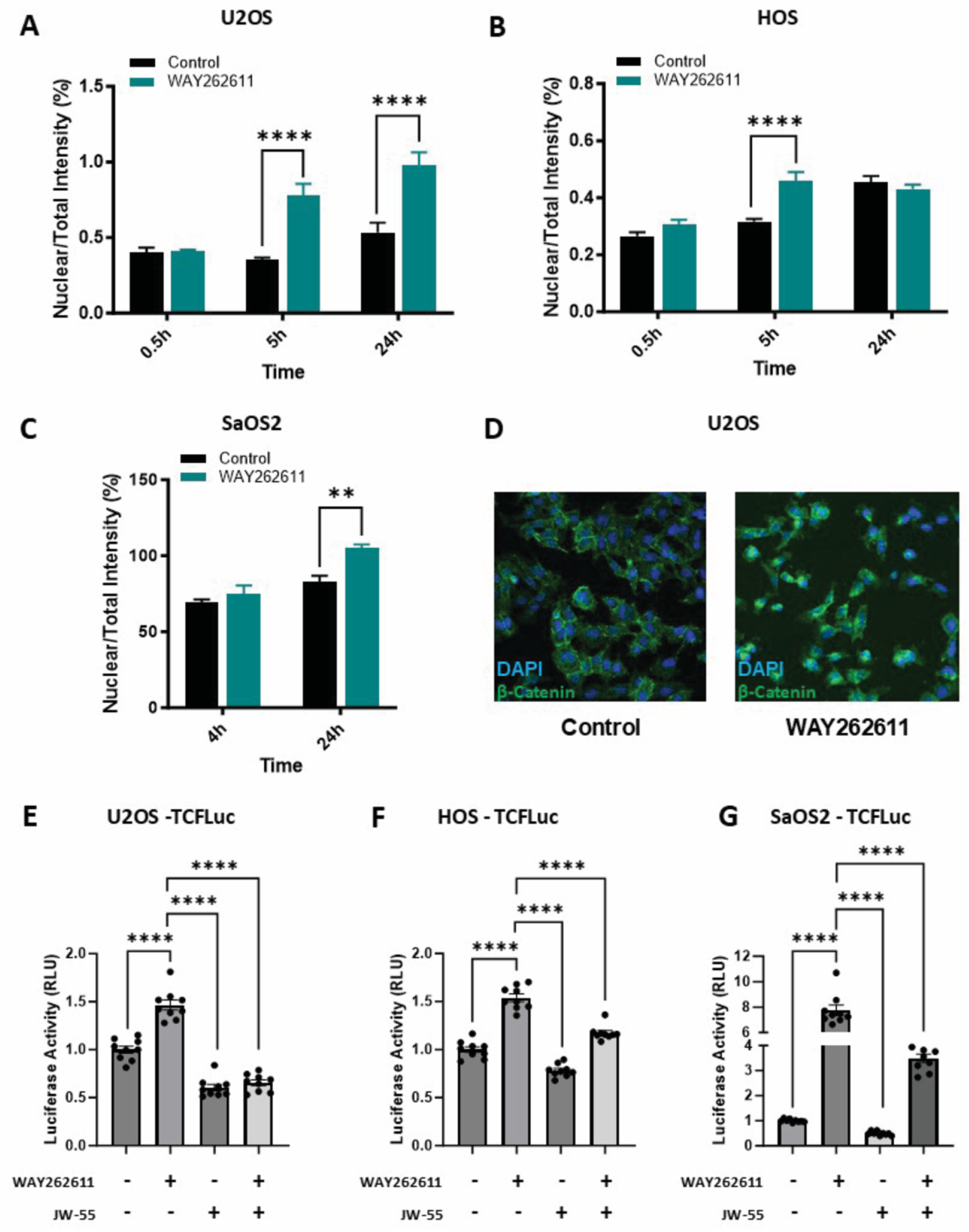
WAY262611 induces nuclear translocation and transcriptional activity of β-catenin. U2OS **(A)**, HOS **(B)**, and SaOS2 **(C)** cells were treated with WAY262611 at the IC50 for each cell line for the indicated duration of time, and β-catenin distribution was evaluated by immunofluorescence. A representative image of U2OS cells is shown in **D**. U2OS **(E)**, HOS **(F)**, and SaOS2 **(G)** cells were stably transfected with a TCF-luciferase reporter construct. Stably transfected cells were treated with either WAY262611 (at the IC50 for each cell line – 3 uM for U2OS, 0.5 uM for HOS and 10 uM for SaOS2), JW-55 (4 uM), or both. WAY262611 induced β-catenin-dependent luciferase activity, and this increase was inhibited by the tankyrase inhibitor, JW-55. JW-55 also attenuated baseline luciferase activity. Error bars are the SEM of triplicate experiments, and each experiment was repeated at least 3 times.

### Induction of Wnt signaling by WAY262611 is dependent on DKK-1

To ensure that the effects of WAY262611 on OS cell lines are due to inhibition of DKK-1 and not off-target effects, we used the CRISPR/Cas 9 gene editing system to knock out DKK-1 in the SaOS2 cell line. The SaOS2 cell line was chosen as a representative OS cell line moving forward. The other cell lines showed decreased DKK-1 with this method, but not a complete knockout. This is not unusual given the vast heterogeneity amongst OS tumors, and is likely representative of the osteosarcoma patient population. Individual clones were generated using flow cytometry assisted cell sorting followed by monoclonal expansion. Knockout of DKK-1 was confirmed by western blot analysis (Figure 4A). A cell line which was transfected with the CRISPR construct but without knockout after monoclonal expansion was used as a control. SaOS2 cells with and without successful knockout of DKK-1 were treated with 6 uM WAY262611 and nuclear localization of beta catenin was analyzed. Unlike control cell lines, the KO cell lines did not have increased beta catenin nuclear localization in response to WAY262611 (Figure 4B). As a functional readout, we investigated the ability of WAY262611 to impact the transcription of genes that are regulated by canonical Wnt signaling in osteosarcoma cells. We found that treating wild type SaOS2 cells with 6 uM WAY262611 resulted in a 75% decrease in expression of ROR1 mRNA, but the drug had no effect on ROR1 expression in the DKK-1 knock out cells (Figure 4C). Taken together, these data support the contention that response to treatment with WAY262611 requires the presence of DKK-1.

**Figure 4.**
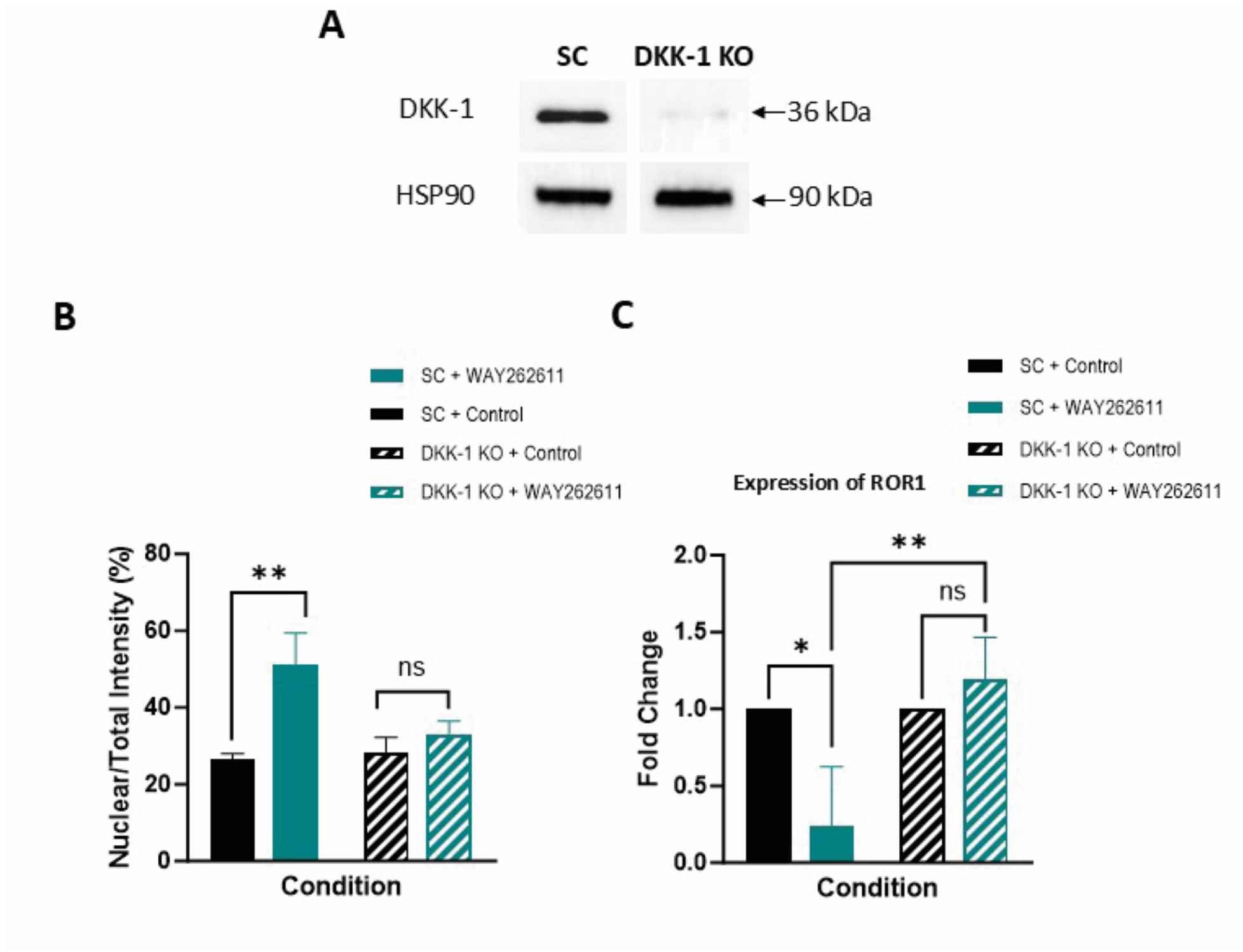
The effect of WAY262611 depends on DKK-1 expression. **(A)** Western blot demonstrating that CRISPR/Cas9 DKK-1 knock-out SaOS2 cells (KO) do not express DKK-1 protein. **(B)** WAY262611 increases nuclear localization of β-catenin in scramble control (SC) cells, but not in KO cells. **(C)** WAY262611 suppresses ROR1 expression in control cells, but not in KO cells. Error bar represents SEM of experiments performed in triplicate. All experiments were performed at least 3 times.

### Treatment with WAY262611 decreases osteosarcoma metastases in a clinically relevant mouse model

Because we previously demonstrated that a neutralizing antibody against human DKK-1 inhibits osteosarcoma metastasis in our clinically relevant mouse model, we next investigated the effect of WAY262611 *in vivo*. There are no published dosing strategies for subcutaneous administration of WAY262611 in mice, so prior to testing the effect of this drug in tumor-bearing animals, we first performed a “mini-Phase I” study to determine the maximally tolerated dose (MTD) of WAY262611 administered by subcutaneous injections in NSG mice. Cohorts of 4 mice were treated with vehicle control (DMSO) or increasing doses of WAY262611. Mice were treated and monitored by body condition score daily for ten days to determine a tolerable dose. Out of all the mice treated with WAY262611, 2 of the mice at the 8mg/kg dose died after developing lethargy and respiratory depression, while the remaining cohorts, including vehicle control, had adequate body condition scores of three and remained healthy and active (Supplemental Figure 1A). Thus, we determined the MTD of WAY262611 to be 4 mg/kg/day in NSG mice. Despite this finding, a preliminary experiment dosing OS PDX-bearing NSG mice at this level resulted in decreased body condition scores, so in further *in vivo* experiments a dose of 2 mg/kg/day was utilized.

To determine the effect of WAY262611 on OS growth and metastasis *in vivo*, we utilized our orthotopic implantation/amputation model.[23] Fragments of an OS PDX were surgically implanted into the tibias of NSG mice. Upon verification of tumor engraftment, mice were randomly assigned to one of 3 cohorts. Cohort A was a control group, treated with DMSO. Cohort B received 2 mg/kg/day of WAY262611 beginning when the tumor was 7 mm in diameter and continued daily until they died or were euthanized. Cohort C received 2 mg/kg/day of WAY262611 after amputation of the tumor-bearing limb once the primary tumor reached 15 mm in maximal diameter and treatment continued daily until they died or were euthanized. Of note, during the establishment of this model, several dozen mice were euthanized at the time of amputation, and necropsy never revealed visually evident metastatic disease. Mice were followed until death or until they were euthanized due to worsening body condition scores (Figure 5A). As was seen using the monoclonal antibody, we found that WAY262611 slows the growth of the primary tumor, resulting in a longer time to amputation (Figure 5B and Supplemental Figure 1B).

**Figure 5.**
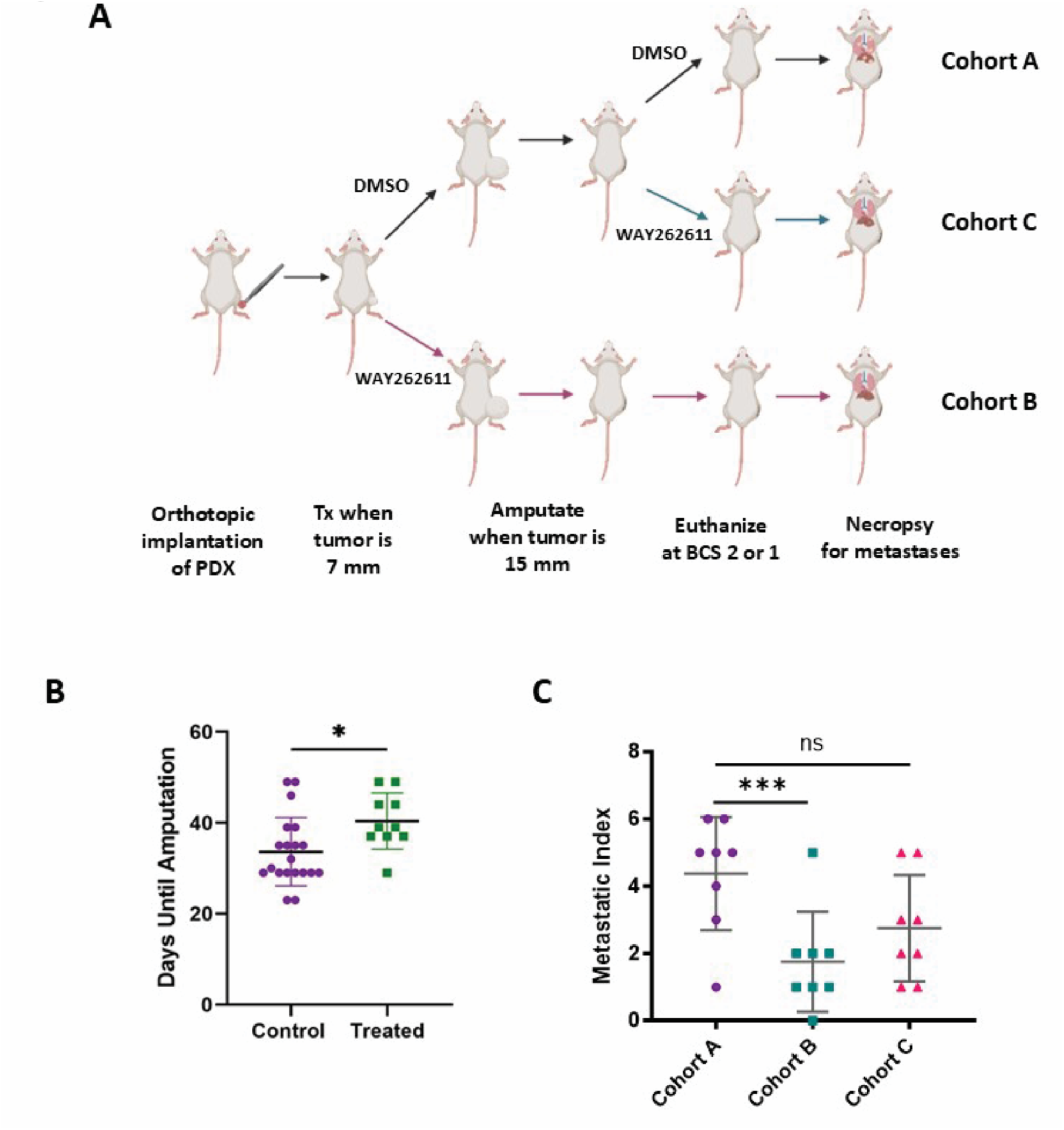
WAY262611 slows the growth of an orthotopically implanted osteosarcoma patient-derived xenograft and inhibits distant metastasis. **(A)** Schematic of the experimental design. (B) Days to amputation of mice treated with either DMSO vehicle control or WAY262611 prior to amputation (Control combines Cohorts A and C). C) WAY262611 reduces metastatic burden, as quantified by metastatic index, when administered pre-amputation, but not when administered post-amputation.

Our laboratory uses a Metastatic Index to quantify the extent of distant hematogenous metastasis in our preclinical experiments. Metastatic lesions were assigned a numeric value according to the size of each tumor (small tumors assigned a value of 1, medium tumors a value of 2, and large tumors a value of 3). For each mouse, the sum of their assigned tumor values was calculated and termed their Metastatic Index. For example, a mouse with 2 small lesions would be assigned a Metastatic Index of 2, whereas a mouse with 1 small lesion and 2 large lesions would be assigned a Metastatic Index of 7. This approach was designed to mimic the standardized sum of cross products utilized to quantify disease response in pediatric oncology clinical trials. The mean Metastatic Index in the control mice was 3.33 ± 0.527 (SEM), compared with 1.00 ± 0.463 in the WAY262611-treated mice prior to amputation (Cohort B). This difference was statistically significant (p=0.005; Figure 5C). The mean Metastatic Index in the mice treated post-amputation was 2.11 ± 0.611, which was not statistically different from control (p=0.067; Figure 5C) but trended strongly in that direction. Thus, mice treated with WAY262611 experienced less metastatic burden than untreated, most prominently in mice treated prior to amputation of the primary tumor, but also in mice treated after removal of the primary tumor. Treatment with WAY262611 had no significant effect on overall survival amongst these cohorts (Supplemental Figure 1C). A photograph of a small metastasis (Supplemental Figure 1D), a large metastasis (Supplemental Figure 1E) and a photomicrograph showing a micrometastasis in the lung (Supplemental Figure 1F) are presented for illustrative purposes.

### WAY262611 induces osteoblastic differentiation

Canonical Wnt signaling plays a key role in normal osteoblastic differentiation. We hypothesized that WAY262611 inhibits osteosarcoma metastasis at least in part because it induces osteoblastic differentiation of osteosarcoma tumors through activation of canonical Wnt signaling, and cancer cells with a more differentiated phenotype are less likely to metastasize. To begin to test this hypothesis, we grew osteosarcoma cell lines in an osteogenic differentiation medium with or without WAY262611 and assessed the expression of two commonly used markers of osteogenic differentiation, alkaline phosphatase (ALPL) and osteopontin (SPP1) by quantitative RT-PCR. We used normal human osteoblasts as a control. Cells were plated on Day-1 in regular growth medium with or without WAY262611 at the IC50 and RNA was harvested the next day to assess the effect of drug alone. On Day 0, cells were plated in differentiation medium, again with or without WAY262611 supplementation. RNA was collected from OS cells on Day 6 and from osteoblasts on Days 3 and 10. As anticipated, human osteoblasts cultured in osteogenic differentiation medium (no drug) expressed steadily increasing levels of both ALPL and SPP1 (Figure 6A). We saw a similar pattern for SPP1 in our osteosarcoma cell lines, with slightly elevated levels initially induced by drug alone, and markedly elevated levels by Day 6 (Figure 6B). The picture with ALPL was less clear. ALPL levels were elevated initially in HOS cells, but were markedly diminished after 6 days in differentiation medium. In contrast, ALPL levels were elevated in U2OS cells but depressed in SaOS2 cells (Figure 6C).

**Figure 6.**
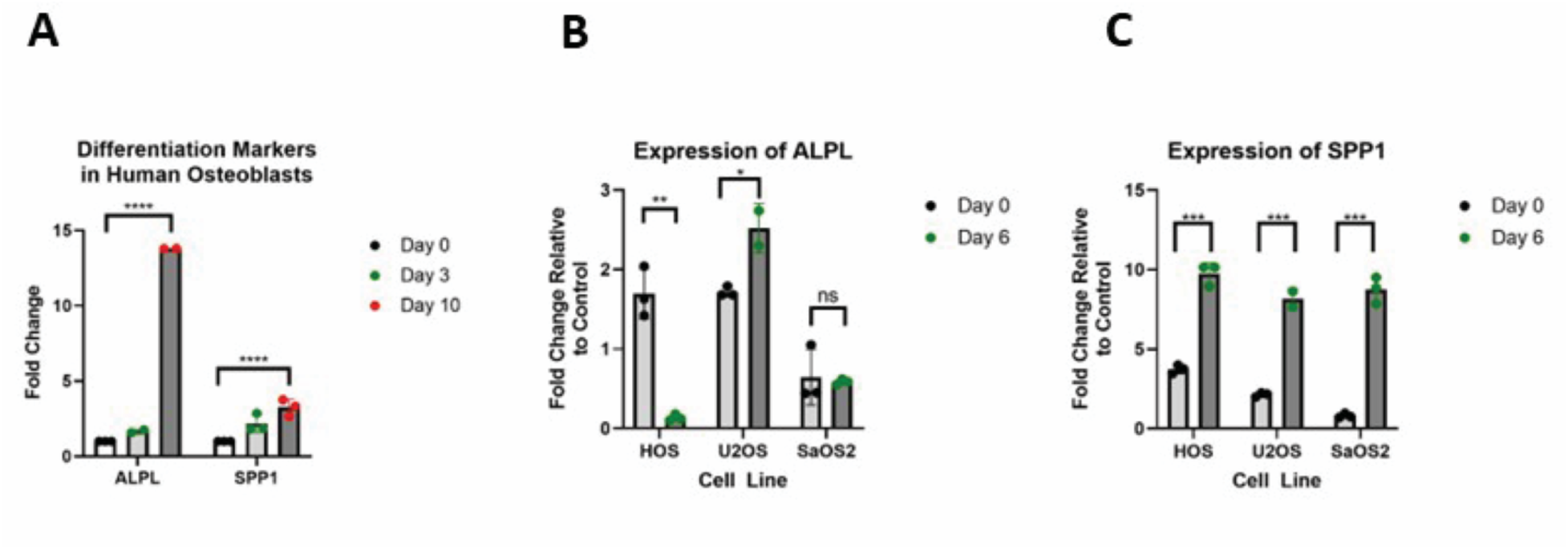
WAY262611 induces expression of molecular markers of osteoblastic differentiation in osteosarcoma cell lines. **(A)** Primary human osteoblasts were cultured in Osteoblast mineralization medium and then supplemented with WAY262611. RNA was harvested on the indicated days and osteopontin (SPP1 and alkaline phosphatase (ALPL) expression were quantified by qRT-PCR. Osteosarcoma cells were cultured in Osteoblast mineralization medium and then supplemented with WAY262611. RNA was harvested on Day 0 (prior to addition of drug) and on Day 6 of drug treatment. SPP1**(B)** and ALPL**(C)** expression was quantified by qRT-PCR. Trend in Panel A statistically analyzed by two way ANOVA. Differences in Panels B and C analyzed by Student’s t Test.

These experiments yielded mixed results and are complicated by multiple confounding issues, including the use of cell lines that have been in *ex vivo* culture for decades and the possibility that ALPL and SPP1 may not be optimal markers of osteogenic differentiation. To better test our hypothesis that WAY262611 induces osteogenic differentiation of osteosarcoma tumors, we performed single cell RNA sequencing (scRNA-seq) of the DAR PDX grown in NSG mice with or without WAY262611 treatment and compared our transcriptomic data with a similar analysis of primary human osteoblasts. Cells isolated from treated tumors resemble each other more than they resemble the untreated tumors (Figure 7A-B). WAY262611-treated tumors show significant enrichment in pathways including DNA methylation, histone lysine demethylation, histone deacetylation, and other epigenetic pathways, mRNA processing, negative regulation of telomere maintenance, and IL-7 signaling (Figure 7C and Supplemental Table 1). Wnt signaling (providing pharmacodynamic confirmation of the mechanism of action of WAY262611) and multiple pathways associated with the extracellular matrix were significantly enriched in the treated cells (Figure 7C). To understand the developmental status of the cells isolated from treated and untreated tumors, we used a scRNA-seq dataset from primary human osteoblasts. [21] This dataset identified 3 subpopulations of normal osteoblasts, including pre-osteoblasts, mature osteoblasts, and an undetermined, intermediate population (Figure 7D). Using this dataset to train pySingleCellNet, a machine learning-based annotation approach, we found that cells from the untreated tumors contained a mixture of cells classified as each of the three subtypes of osteoblasts. In contrast, the majority of the cells isolated from the WAY262611-treated tumors were more similar to mature osteoblasts (Figure 7E-G). Taken together, these data strongly support our hypothesis that WAY262611 induces a more differentiated phenotype in human osteosarcomas.

**Figure 7.**
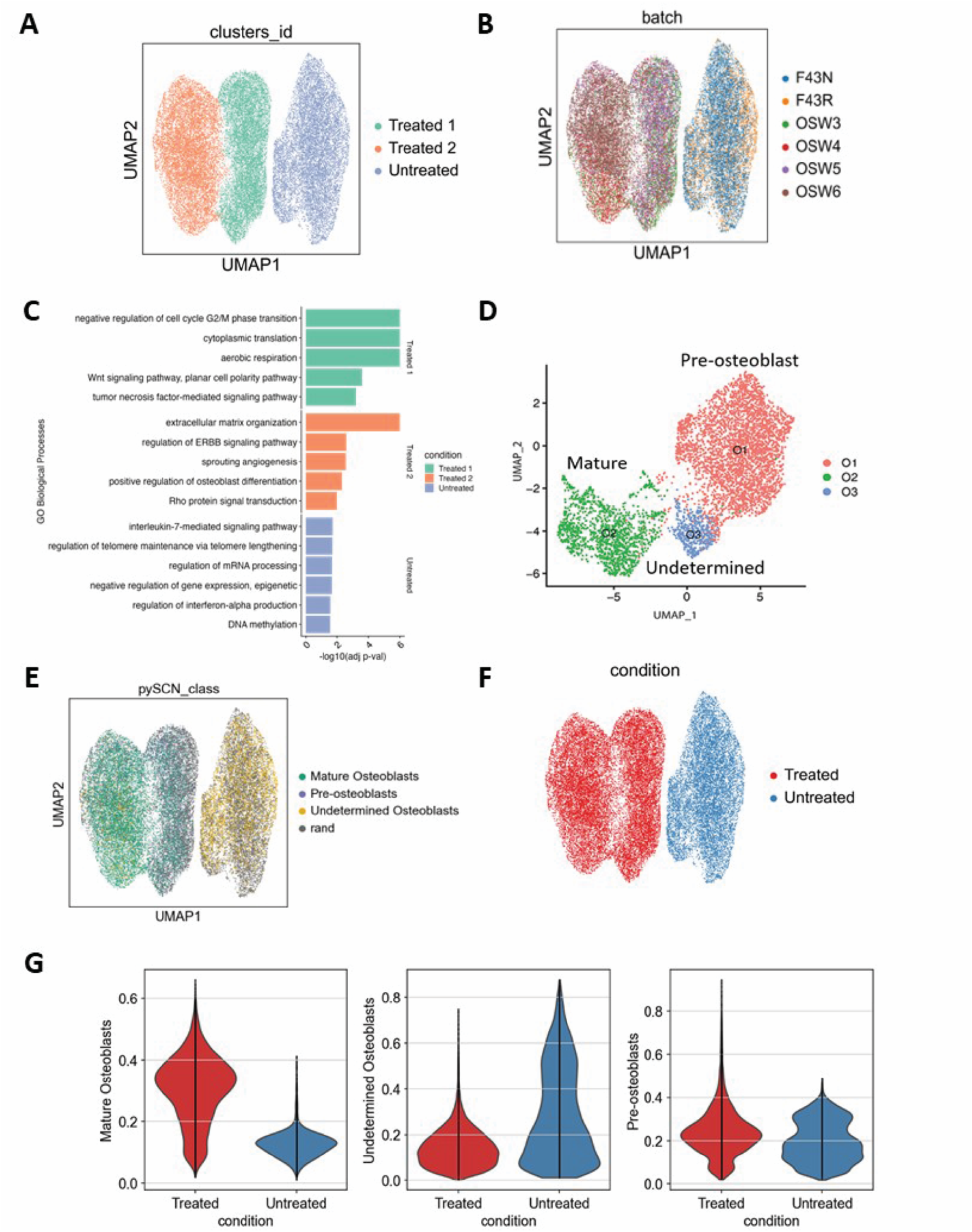
Single cell RNA sequencing reveals that WAY262611-treated osteosarcoma patient-derived xenografts demonstrate increased differentiation. **(A)** UMAP analysis shows that cells isolated from untreated and from treated tumors exist in 3 distinct transcriptional states (the untreated tumors in one state, and the treated tumors in 2 others, with little overlap between treated and untreated). **(B)** Batch effect analysis confirms that the untreated cluster contains cells from the 2 untreated tumors randomly dispersed within the cluster, and that the 2 clusters from treated tumors both contain cells from each of the 4 treated tumors analyzed. **(C)** GSEA reveals a number biological processes enriched in each cell cluster. **(D)** UMAP analysis of primary human osteoblasts reveals 3 distinct transcriptional states, labeled O1, O2, and O3, corresponding to pre-osteoblasts (O1), mature osteoblasts (O2), and an indeterminate state (Undetermined; O3). **(E)** Data from normal human osteoblasts were used to train a pySingleCellNet classifier, and this was used to evaluate the cells isolated from both treated and untreated PDX. The Untreated cluster is composed predominantly of pre-osteoblasts and undetermined osteoblasts, whereas Treated Cluster 1 is composed predominantly of mature osteoblasts and Treated Cluster 2 is predominantly pre-osteoblasts and mature osteoblasts. **(F-G)** Quantification of the data from the pySingleCellNet classifier shows a selective increase in mature osteoblast-like cells in the treated tumors compared to the untreated tumors, with no significant difference in the other two states.

## Discussion

Although the introduction of systemic chemotherapy dramatically improved the survival of patients diagnosed with localized osteosarcoma, there have been no improvements in the survival of patients diagnosed with metastatic disease, nor those who suffer a metastatic recurrence, despite multiple clinical trials testing ever-more-intensive chemotherapy regimens. Improved outcomes for these patients are therefore more likely to result from therapies aimed at the biology of metastasis. To this end, we have been investigating the role of DKK-1, and Wnt signaling in general, in osteosarcoma metastasis. We previously demonstrated that a neutralizing monoclonal antibody against DKK-1 prevents osteosarcoma metastasis in our clinically relevant mouse model. The current work advances that finding by demonstrating that a small molecule inhibitor of DKK-1, WAY262611, activates canonical Wnt signaling in osteosarcoma cell lines *in vitro*, drives the differentiation of osteosarcoma tumor cells *in vivo*, and significantly impairs spontaneous distant metastasis in a mouse model.

There are several important implications of this work. Most importantly, taken in combination with our previous work and those of other laboratories, this report takes a big step toward credentialing DKK-1 as a valid therapeutic target in osteosarcoma. The Gregory laboratory demonstrated that patients with osteosarcoma have elevated levels of DKK-1 in their blood at the time of diagnosis [14], and later demonstrated that DKK-1 upregulates expression of genes involved in treatment resistance, such as ALDH1A1.[24] They also have shown that morpholino-directed downregulation of DKK-1 slows the growth of murine osteosarcoma tumors.[25] Taken altogether, their work and ours provide substantial evidence that targeting DKK-1 can interfere with osteosarcoma tumor growth and metastasis and that this protein represents a valuable potential therapeutic target that should be explored in much greater depth.

In our previous publication, we found that removing DKK-1 from the plasma of tumor-bearing mice with a neutralizing antibody led to increased expression of markers of osteoblastic differentiation in osteosarcoma tumors growing in an orthotopic location. The current work expands upon that finding and strengthens support for the hypothesis that inhibition of DKK-1 promotes the osteoblastic differentiation of osteosarcoma cells, and that this is the mechanism by which this approach inhibits metastasis. *In vitro*, we found that DKK-1 inhibition promotes the expression of SPP1, or osteopontin, a vitamin D3-regulated protein that plays a key role in normal bone mineralization, being expressed in pre-osteoblasts, osteoblasts, and osteocytes, among many other cell types. We performed scRNA-seq analysis of a human osteosarcoma PDX before and after treatment with WAY262611 and compared our results with the dataset published by the Deng laboratory. We found that untreated tumors contain cells most similar to both pre-osteoblasts and to mature osteoblasts, and that treatment with WAY262611 causes a significant shift in the transcriptomes of these cells toward mature osteoblasts. This, along with the enrichment of ECM seen by GSEA, provides strong evidence that inhibition of DKK-1 promotes the differentiation of osteosarcoma cells towards a mature osteoblast-like state and suggest that this treatment approach might represent a new manifestation of so-called “differentiation therapy.”

Over the past 30 years, differentiation therapy has been gaining traction as a non-chemotherapy approach to treat malignancies.[26–28] The best example of differentiation therapy is the use of all-trans retinoic acid to treat acute promyelocytic leukemia.[29, 30] This innovation, along with the addition of a second differentiation agent, arsenic trioxide, increased 2-year overall survival from 20% to over 90% and resulted in a profound shift in the standard of care for this myeloid malignancy such that newly diagnosed patients are no longer treated with cytotoxic chemotherapy. [31] In solid tumors, the most successful application of differentiation therapy is the case of high-risk neuroblastoma, where the addition of cis-retinoic acid, which induces differentiation of neuroblastoma cells *in vitro*, to chemotherapy resulted in a significant improvement in 5-year event-free survival and is now part of the standard of care for children with this aggressive cancer. [32, 33] Future development of DKK-1-targeting agents has the potential for successful combination with standard chemotherapy and may improve outcomes by limiting the risk of metastatic relapse. We envision future in vivo studies focused on combination therapy to understand if there is synergy or benefit to combination therapy. Initial clinical trials will likely be proposed in a multiple metastatic or refractory patient population in combination with chemotherapy to treat active disease and then prevent its further spread. Thereafter, we aim to propose trials using maintenance monotherapy after systemic therapy and surgery for localized osteosarcoma with the goal of preventing metastatic spread in patients without metastases at initial diagnosis. With advancements in the use of circulating tumor DNA, we can use peripheral samples to monitor for relapse in addition to standard imaging techniques in these scenarios.

There are limitations to our study that require further investigation. We treated mice with WAY262611 until death, which is not generally how targetable agents are administered in adjacent clinical scenarios. We are proceeding with studies on a new monoclonal antibody to DKK-1, DKN-01, which is a more clinically relevant formulation. In these future in vivo studies, we plan to have cohorts of mice that will stop the inhibitor at various time points and then be monitored for time to development of metastases. Additionally, the use of an immunocompromised mouse model may not reflect disease response accurately or the adverse effects that patients may manifest when receiving the inhibitor. Given our knowledge about other therapies targeting differentiation, we may expect to see differentiation syndrome, especially since the canonical Wnt pathway not only affects osteoblasts but is omnipresent in critical organ systems. As in other treatments, we are generally able to treat patients with differentiation syndrome with aggressive supportive care.

Although our analysis was focused on the effect of DKK-1 on the metastatic capacity of the primary tumor, it is also possible that inhibition of DKK-1 affects the metastatic niche. Multiple studies have demonstrated that osteosarcoma tumors can influence sites of distant metastases, such as the lungs, through secretion of extracellular vesicles that can induce changes in the lungs of tumor-free mice in ways that would be expected to promote metastatic colonization.[34–38] It is possible that these or other changes that establish a premetastatic niche might be regulated by Wnt signaling, and thus be impacted by treatments that inhibit DKK-1. Another potential mechanism by which DKK-1 might modulate osteosarcoma metastasis is by affecting the functional interactions among early disseminated tumor cells and niche cells. The Roberts laboratory has put forward a model wherein lungs are initially colonized by “anchor cells” that establish an initial micrometastasis through interactions with various populations of lung cells (alveolar macrophages, fibroblasts, and others), and that this initial lesion can then support the proliferation of “growth cells” that drive the development of the macrometastatic lesions. [39] It is possible, though it has not yet been investigated, that these interactions can be modified by treatments targeting the Wnt signaling pathways, and that this can contribute to our observation that inhibition of DKK-1 suppresses the development of visceral osteosarcoma metastasis. Another potential explanation for our findings is an effect of WAY262611 on the ability of disseminated tumor cells to home to secondary sites of disease, a mechanism that will also be further explored in the future.

In summary, we found that inhibition of DKK-1 by the small molecule WAY262611 activates canonical Wnt signaling, promoting the differentiation of osteosarcoma both *in vitro* and *in vivo*, and that this results in decreased visceral metastasis in our clinically relevant mouse model. This approach is readily translatable, as multiple pharmaceutical companies have developed antibodies that target DKK-1 for therapeutic use. This work, therefore, suggests a potential path forward in the prevention of osteosarcoma metastasis, whether derived from a primary tumor or from established metastases. This would be a significant advance in the treatment of high-risk osteosarcoma, a tumor that has seen almost no therapeutic improvement in decades.

## Acknowledgements

This work was supported by grants from the NIH (R01CA262802 to DML and PC, U19AG055373-08 and 5P20GM109036-08 to H-WD) and the Montefiore Einstein Comprehensive Cancer Center Support Grant 2P30CA013330-5.

## Notes

The authors declare no potential conflicts of interest

### Competing Interest Statement

The authors have declared no competing interest.

