## Supplemental Figure 1 for "Inhibition of DKK-1 by WAY262611 Inhibits Osteosarcoma Metastasis"

**Supplemental Data**

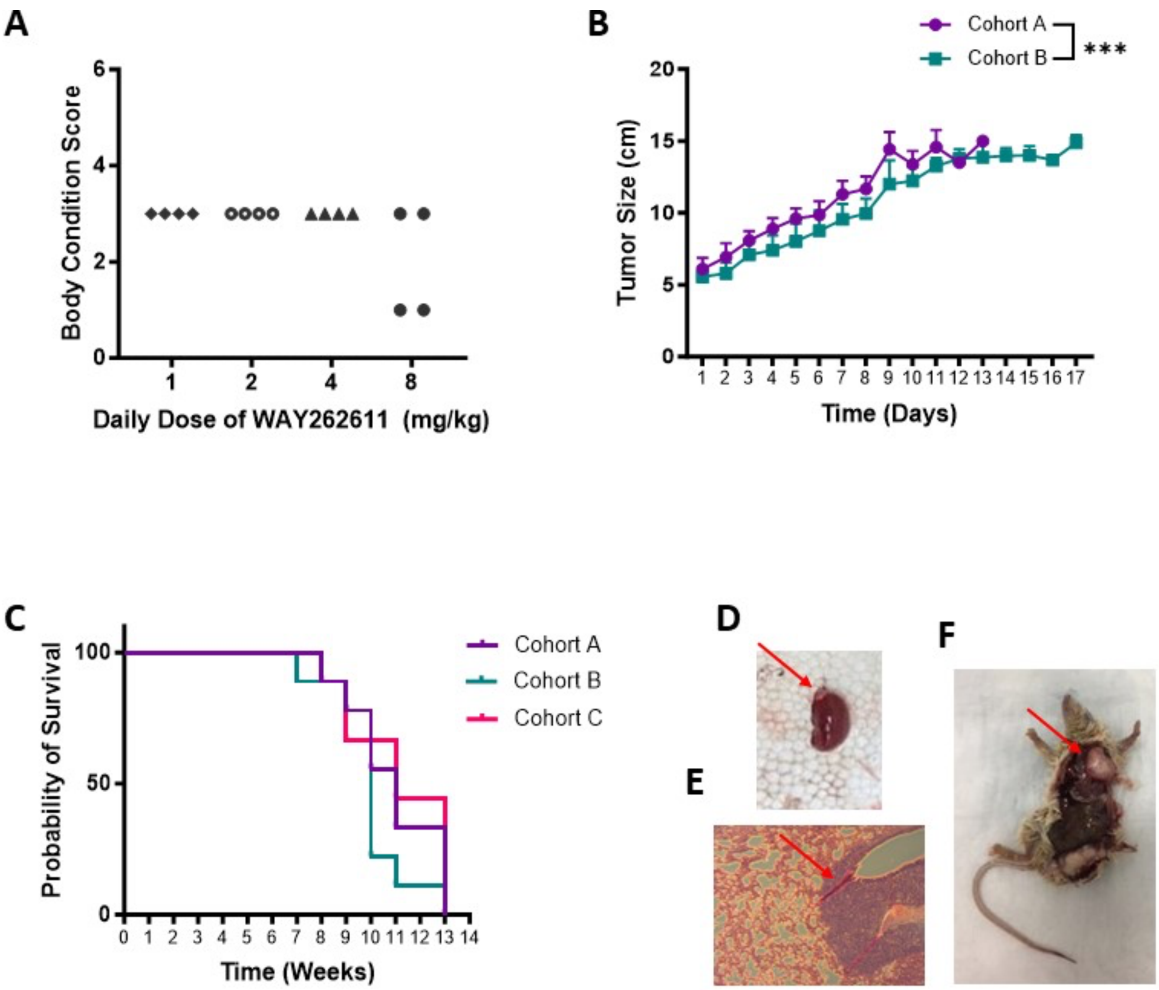


**Supplementary Figure 1.**

WAY262611 slows the growth of orthotopically implanted osteosarcoma PDX and inhibits metastasis, but does not affect survival. We used a "mini-Phase I" study to identify an appropriate does of WAY262611 in NSG mice. **(A)** Body Condition Score only decreased in mice treated for 10 days with 8 mg/kg/day WAY262611 in DMSO. Each symbol represents an individual mouse. **(B)** Tumor fragments from the DAR PDX were implanted orthotopically in the pretibial space, and tumor size was measured in mice treated with vehicle control (DMSO, Cohort A) or WAY262611 (Cohort B). There were 5 mice per cohort, and the error bars represent the SEM. The tumors in the treated mice grew more slowly than those in control mice. These curves are statistically significantly different by 2-way ANOVA ($p<0.001$). **(C)** Survival curve of control mice (DMSO, Cohort A), mice treated with WAY262611 from the time of tumor engraftment (Cohort B), and mice treated after amputation (Cohort C). Treated mice did not have improved survival. **(D)** The red arrow indicates a small renal metastasis. **(E)** Photomicrograph of a histologic section through the lungs of a mouse from Cohort A. The red arrow indicates a focus of metastatic osteosarcoma. **(F)** A large pulmonary nodule seen at necropsy is indicated by the red arrow.
